# The power of dying slowly - persistence as unintentional dormancy

**DOI:** 10.1101/2021.01.20.427471

**Authors:** João S. Rebelo, Célia P. F. Domingues, Francisca Monteiro, Teresa Nogueira, Francisco Dionisio

## Abstract

Persistence is a state of bacterial dormancy where cells with low metabolic activity and growth rates are phenotypically tolerant to antibiotics and other cytotoxic substances. Given its obvious advantage to bacteria, several researchers have been looking for the genetic mechanism behind persistence. However, other authors argue that there is no such mechanism and that persistence results from inadvertent cell errors. In this case, the persistent population should decay according to a power-law with a particular exponent of −2. Studying persisters’ decay is, therefore, a valuable way to understand persistence. Here we simulated the fate of susceptible cells in laboratory experiments in the context of indirect resistance. Eventually, under indirect resistance, detoxifying drug-resistant cells save the persister cells that leave the dormant state and resume growth. The simulations presented here show that, by assuming a power-law decline, the exponent is close to −2, which is the expected value if persistence results from unintentional errors. Whether persisters are cells in a moribund state or, on the contrary, result from a genetic program, should impact the research of anti-persistent drugs.

**Author Summary:** Persistence, a form of bacterial dormancy, was discovered in the early days of the antibiotic era. Thanks to dormancy, these cells often evade antibiotic therapy and the immune system. However, despite its clinical importance, this phenotype’s nature is still under debate. Arguably, the prevailing view is that persistence is an evolved (selected for) bet-hedging mechanism to survive in the presence of cytotoxic agents such as antibiotics. In that case, the persister population should decay exponentially, although at a much slower pace than the non-persister population. A few authors recently advanced an alternative hypothesis: bacterial persistence results from many malfunctions and cell division errors. In this case, persistent populations should decay according to a power-law with exponent of −2, that is, according to 1/t^2^. Here we simulated the fate of susceptible bacterial cells in the presence of bactericidal antibiotics in the context of indirect resistance based on laboratory experiments performed earlier. By showing that the dynamics of persister cells is consistent with 1/t^2^, our results corroborate the hypothesis that the phenomenon of bacterial persistence is an accidental consequence of inadvertent cell problems and errors. If confirmed, this conclusion should impact the research strategies of anti-persistent drugs.

“The following day, no one died. This fact, being absolutely contrary to life’s rules, provoked enormous and, in the circumstances, perfectly justifiable anxiety in people’s minds, for we have only to consider that in the entire forty volumes of universal history there is no mention, not even one exemplary case, of such a phenomenon ever having occurred…”*Death with interruptions*
José Saramago (2005)
Nobel Prize for Literature 1998

## Introduction

Susceptible bacterial populations do not perish instantaneously in the presence of bactericidal antibiotics. Instead, they decay exponentially, typically for a few hours for wild-type bacterial strains. After this first period, a second population has a significantly lower death rate [1,2]. These cells are in the persistent state and usually account for less than 1% of the original bacterial community. The persistent cells do not harbor resistance genes, but they thrive in the presence of a drug or other harsh environments by lowering their metabolic activity and growth rate. Importantly, because of their ability to resume growth following antibiotic therapy, they are responsible for recurrent and chronic infections [2–4]. The ubiquitous distribution of persistence among bacteria, fungi, and cancer cells [2,5,6], together with its impact on antibiotic resistance development among bacteria [7,8], highlights the need for a better understanding of the role of bacterial persistence in the survival of pathogenic bacteria.

Persistence is often involved in indirect resistance [9]. During indirect resistance, susceptible cells are protected against a bactericidal antibiotic because other co-inhabiting bacterial cells detoxify the medium through antibiotic degradation or modification [10,11]. Once the environment becomes nontoxic, cells that leave the persistent state survive and thrive [9]. Indirect pathogenicity is an alternative name for indirect resistance because, in many cases, it involves antibiotic-susceptible pathogenic bacteria and cells from a non-pathogenic bacterial species that detoxify the environment enabling the growth of the pathogens (see, for example, refs. [10,12,13].

In the context of indirect resistance, the chances of survival of susceptible cells depend on several factors, not just entering into the dormant state of persistence. For example, medium detoxification certainly takes some time to be completed. Moreover, the survival of susceptible cells depends on detoxifying cells’ density and the total cell density. High population density implies that susceptible and resistant cells tend to be close neighbors, increasing the odds of susceptible cells [14–16]. Furthermore, the survival of susceptible cells should depend on the death rate of non-persister cells in the presence of antibiotics and, importantly, on the persistent cells’ behavior and death rate.

While the bactericidal antibiotic is still present, any bacterium returning to growth dies. As mentioned above, in the presence of a bactericidal antibiotic, the non-persistent population of wild-type strains declines exponentially (that is, according to exp(−k.t), where t is time, and k is a constant). The exponential decay is a direct consequence of the fact that bacterial clonal populations are homogeneous and involve many independent entities (cells), each having the same constant probability per unit of time of starting growth. After some time of decaying exponentially and fast, a second phase begins. In this phase, only persister cells are alive. This population also decays, although at a lower rate [1], because persister cells die if they return to growth while the medium is still toxic.

Until recently, the assumption was that the persistent population also decays exponentially, but at a much slower pace. However, some studies have suggested that the persistence state results from different kinds of faults and errors in cell division rather than an evolved genetic program [17–20]. If true, the persistent population is physiologically heterogeneous, comprising several sub-populations, each with its proper exponential decay. The sum of all these negative exponentials results in a power-law curve (i.e., proportional to t^α^, where t is time and α is the exponent), instead of exponential decay [21]. Importantly, this reasoning also tells us that the exponent of the power-law decay should be close to −2, which was experimentally corroborated [21]. In other words, the population of persister cells should decay proportionally to t^−2^, where t is time. However, there is no way to calculate the rate of exponential decay (that is, there is no way to compute the value of the constant k in exp(−k.t)).

Fig 1 shows what happens to a clonal drug-sensitive bacterial population exposed to a cytotoxic drug, as well as the importance of a power-law decay. In both Figs 1a and 1b, the first decaying phase consists of exponential decay but, after some time (instant t = τ_o_), the decline is much slower. However, even if the exponential decay is prolonged, it crosses the power-law decay sooner or later (Fig 1b). This fact’s biological meaning is that a power-law decline may allow the persistent population’s survival for much longer. The long tail of power-law distributions is relevant for indirect resistance because resistant (detoxifying) cells may take a long time to detoxify the medium.

**Fig 1.**
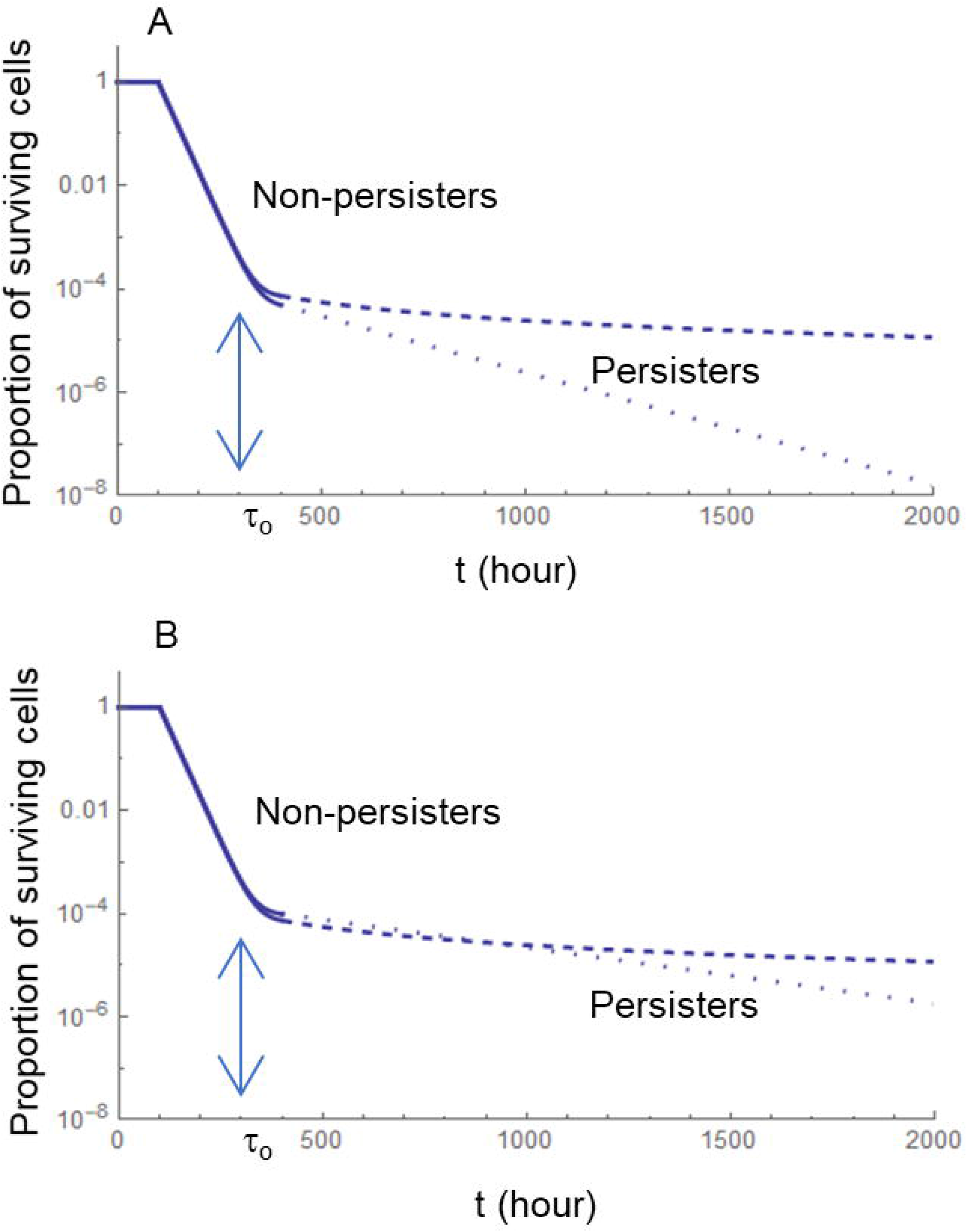
Decay of a drug-susceptible bacterial population in the presence of a bactericidal antibiotic. The horizontal axis, representing time t (hours), is linear, while the vertical axis is on a logarithmic scale and represents the proportion of the population that is still alive. Descending full lines represent the exponential decay of the non-persistent population, according to exp(−0.04 t). When t = τ_0_, only persisters are alive. The broken lines represent decay according to the power-law 1/t^2^. The dotted lines represent exponential decay, i.e., according to exp(−k t). A: persisters decay according to constant k = 0.005 h^−1^. B: persisters decay according to constant k = 0.0025 h^−1^.

This paper’s general goal is to investigate the behavior of persistent populations. To achieve this goal, we studied the behavior of genetically susceptible cells in the context of indirect resistance. Therefore, we had two main objectives. First, we aimed to understand whether persistence is necessary for the survival of sensitive cells in the context of indirect resistance (Fig 2). Second, we aimed at understanding whether the decay of the persistent population is better explained by a negative exponential or by power-law with a negative exponent. To achieve these two main objectives, we took advantage of previous experiments performed in our laboratory, where we measured the degree of protection of susceptible cells when co-cultured together with β-lactamase-producing cells and in the presence of the β-lactam antibiotic ampicillin [14]. In the present paper, we performed computer simulations to understand how many non-persister and persister cells contributed to the survival of susceptible cells, identified which parameters explain such contributions, and directly compared results with those obtained experimentally by [14]. In the end, we hoped to understand persistent cell formation mechanisms, which is critical for developing medical strategies against pathogenic bacterial persisters.

**Fig 2.**
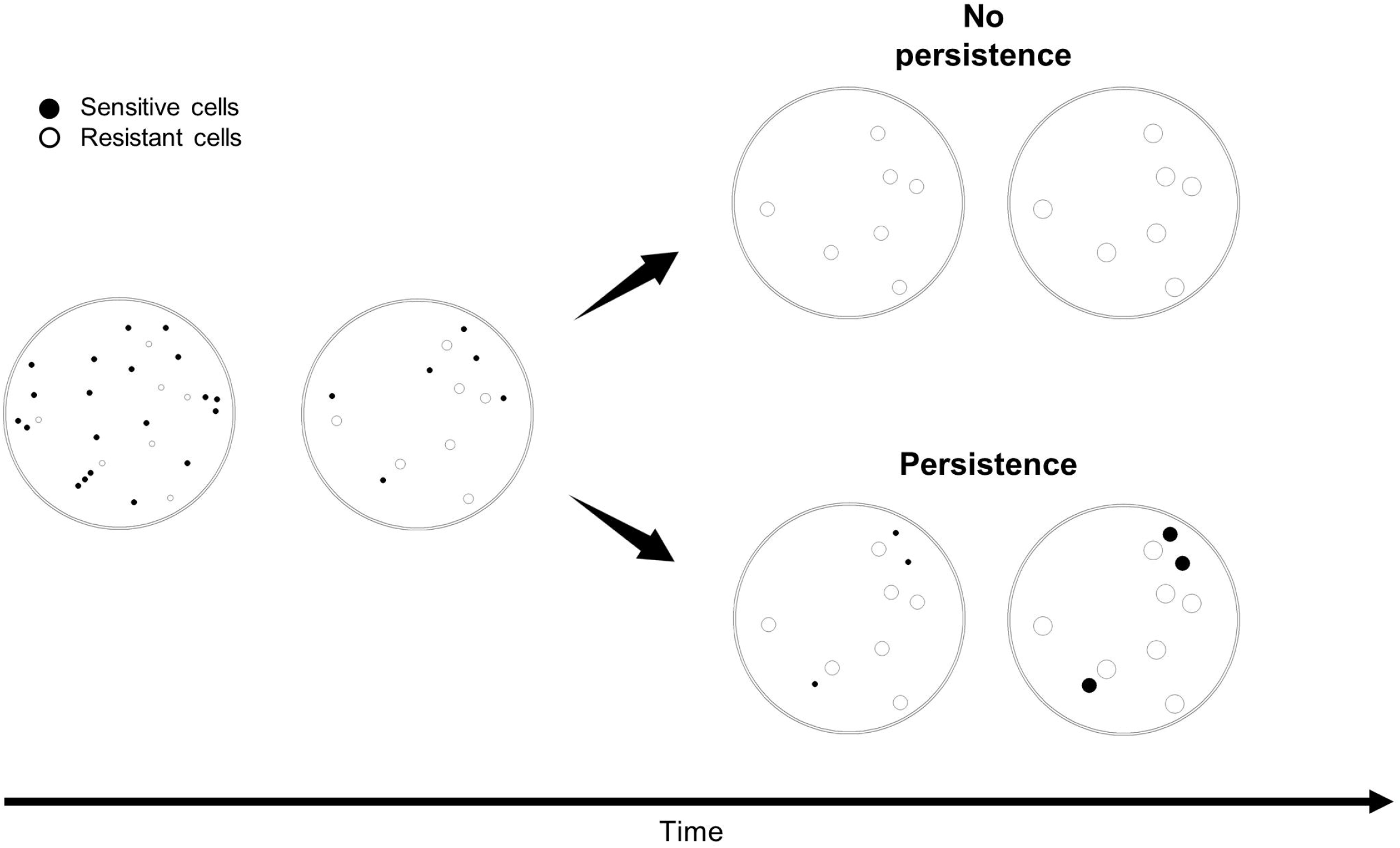
Indirect resistance and the survival of susceptible cells through persistence. When exposed to antibiotics, persister cells survive and eventually grow after medium detoxification by resistant cells. Blue circles represent susceptible cells, and orange circles represent resistant cells.

## 2. Methods

### Previous experimental data used in this study

In the context of indirect resistance, we intend to understand how susceptible bacteria survive while the medium is still toxic. For that, we compared experimental results from our previous experiments [14] with simulations performed in the present work. In those experiments, we used bacterial cells of *Escherichia coli* to measure the degree of protection of susceptible cells in the presence of ampicillin when co-cultured with cells encoding a β-lactamase (resistant cells). The basic experimental setup was to initiate the co-culture with a specific total initial density and frequency of resistant:sensitive cells in plates with rich medium supplemented with ampicillin. After incubating for 24h, we quantified the density of both susceptible and resistant cells. These cells were resistant to ampicillin because they harbored the natural isolated R1 plasmid, which encodes a β-lactamase that detoxifies the medium by breaking the β-lactam ring through hydrolyzation. This plasmid is conjugative, so we also quantified transconjugants (here defined as cells that received the plasmid plus their descendants). However, the frequency of transconjugants remained very low [14], which is a consequence of the fact that the conjugation rate of the R1 plasmid in the *E. coli* strain used in the experiments is low [22–24].

To develop our study, we used the experimental data for (i) two initial total cell densities – approximately 10^7^ cfu/mL and 10^5^ cfu/mL, henceforth denominated as high and low density respectively; and (ii) three proportions between resistant (R) and susceptible (S) cells – 1R:99S, 50R:50S and 99R:1S (where, e.g., 1R:99S means a frequency of a resistant cell for 99 susceptible cells). The relevant information about the initial experimental conditions and final results from the Domingues *et al.* study (ref.[14]) are in Table 1, where we can see the average of three replicates.

**Table 1.** Experimental data (average values) from ref. [14]

| Density | Frequency | Initial resistant | Final resistant | Initial susceptible | Final susceptible | Transconjugants |
| --- | --- | --- | --- | --- | --- | --- |
| Low | 1R:99S | $2.12 \times 10^3$ | $8.87 \times 10^9$ | $2.16 \times 10^5$ | $2.03 \times 10^2$ | 0 |
| | 50R:50S | $1.22 \times 10^4$ | $3.90 \times 10^{10}$ | $1.04 \times 10^4$ | $1.70 \times 10^1$ | 0 |
| | 99R:1S | $1.87 \times 10^5$ | $1.40 \times 10^{10}$ | $2.03 \times 10^3$ | $1.70 \times 10^1$ | 0 |
| High | 1R:99S | $7.00 \times 10^4$ | $2.04 \times 10^{10}$ | $2.93 \times 10^6$ | $3.27 \times 10^4$ | $1.03 \times 10^2$ |
| | 50R:50S | $5.00 \times 10^5$ | $7.70 \times 10^9$ | $4.95 \times 10^5$ | $1.08 \times 10^6$ | $1.60 \times 10^4$ |
| | 99R:1S | $5.40 \times 10^7$ | $6.53 \times 10^9$ | $8.73 \times 10^5$ | $1.23 \times 10^7$ | $1.39 \times 10^5$ |

### Computational Model - flow of the simulation

Here we describe the algorithm of the simulation process. Table 2 and Fig 3 show the respective pseudocode and flowchart. All code is available on GitHub (https://github.com/jrebelo27/Simulation-code-of-persistence).

**Table 2.**
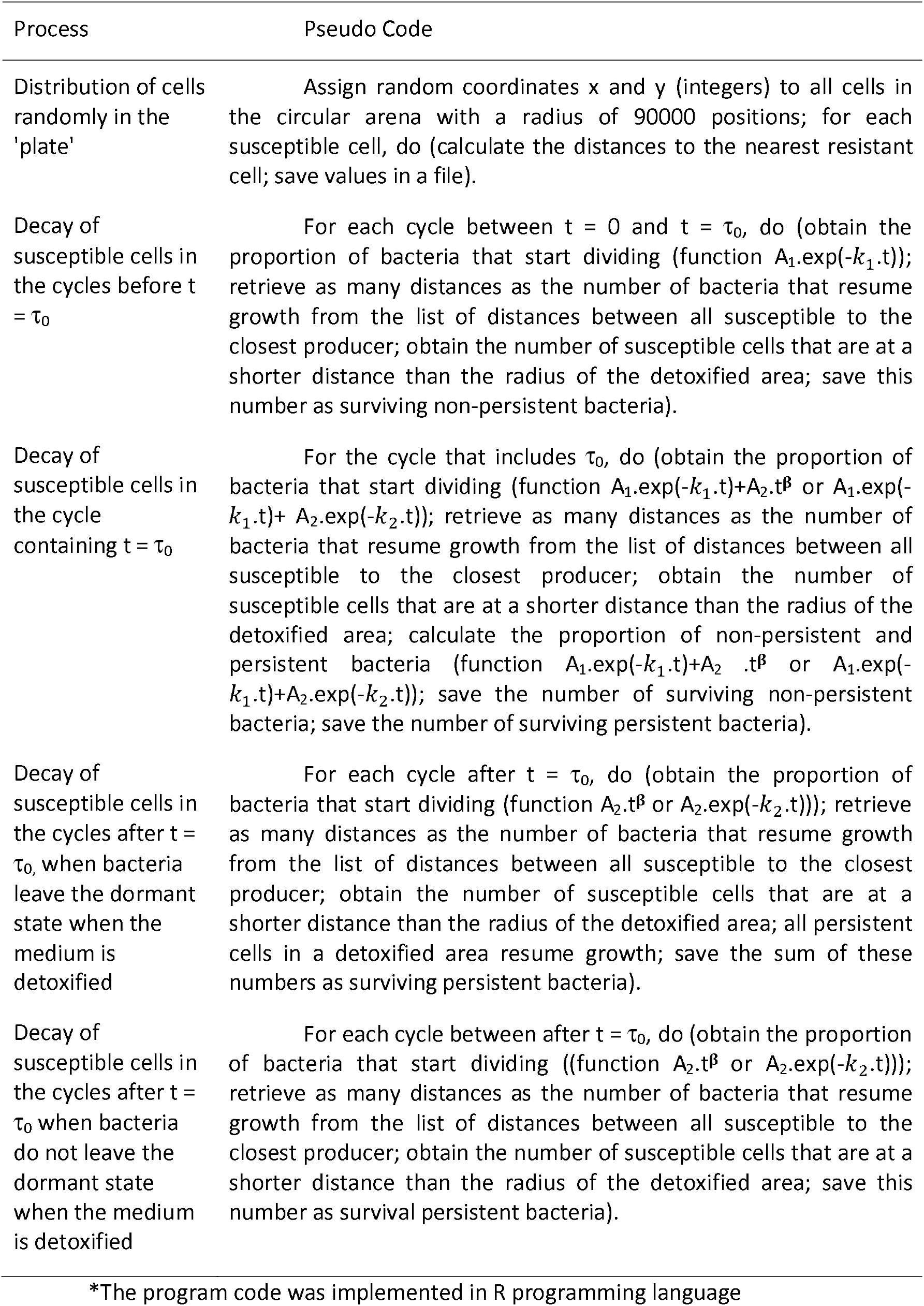
Pseudocode of the program*

**Fig 3.**
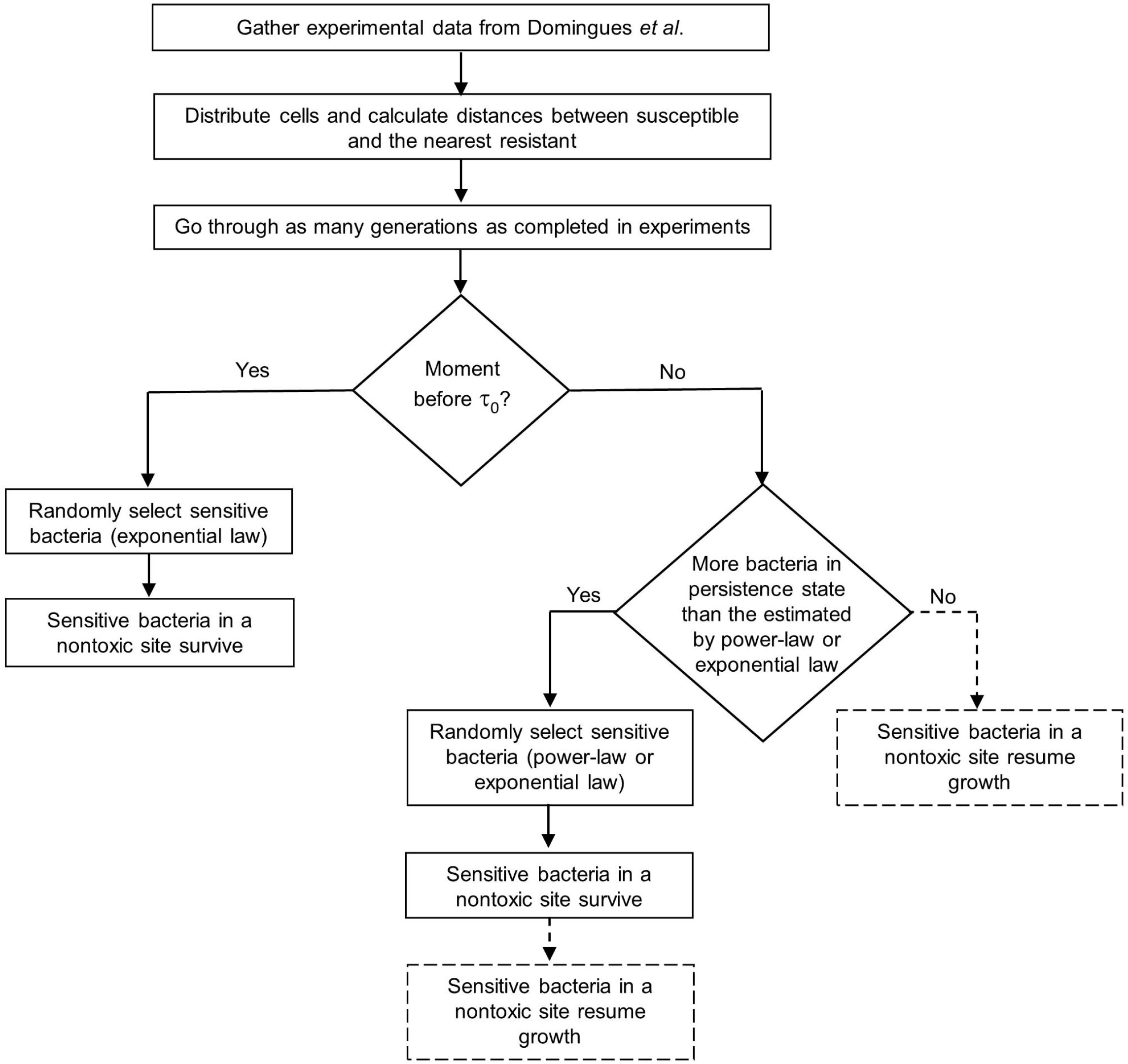
Flowchart of the program. After distributing cells in the ‘plate’, the program simulates bacterial growth during as many generations as the ones completed in experiments of ref. [14]. The decay of the bacteria varies depending on the time interval in which the simulation is. Dotted lines only happen when the biological assumption is that persister cells leave the dormant state as soon as their site is nontoxic.

We simulated the spread of resistant and sensitive cells with a given total density and at specific proportions (1R:99S, 50R:50S, or 99R:1S) in a medium plate, and we saved in a file the distances between each susceptible cell and the nearest resistant cell.

As in the experiments by Domingues *et al.* (ref. [14]), we assumed that the plate contains nutrients and the antibiotic ampicillin. Susceptible cells are, by definition, sensitive to this antibiotic, and resistant cells can detoxify their surroundings, clearing up the cytotoxic antibiotic. A decreasing ampicillin concentration gradient is generated from the inside out by the diffusion of the β-lactamase enzyme that degraded the antibiotic.

The simulation is composed of several cycles, as much as the number of generations completed by the resistant cells in the experiments by Domingues *et al.* (ref. [14]). The following happens in each cycle: there is detoxification of a specific circular area around every resistant cell, simulating the spread of β-lactamase. Such spread occurs for a certain time, the equivalent of a bacterial generation. Meanwhile, resistant cells replicate once.

Regarding sensitive bacteria, non-persistent and persistent behave differently:

i. Non-persister cells: the computer program randomly takes a percentage of susceptible bacteria, defined according to the exponential decrease (Fig 1), and tests whether its site is already nontoxic (that is, if the distance to the nearest resistant cell is less than the total detoxified radius around the resistant cell). If yes, the susceptible bacterium survives. If not, that susceptible cell dies; in practice, the program removes that cell from the simulation’s next steps.
ii. For persister cells, the computer program may follow two different approaches, depending on the biological assumptions. Either (a) persister cells leave the dormant state stochastically only according to the function considered (exponential-law or power-law), or (b) persister cells resume growth (leaving the dormant state) stochastically according to an exponential or power-law distribution or whenever their site becomes detoxified. If the biological assumption is that persister cells leave the dormant state only stochastically (independently of the antibiotic’s presence/absence), the computer program follows the approach (a). In this case, a percentage of susceptible bacteria is randomly taken (according to the power-law or the exponential-law) (Fig 1). The program tests whether each susceptible cell’s site is already nontoxic (that is, if the distance of each susceptible (persister) cell to the nearest resistant cell is less than the total detoxified radius around the resistant cell). If yes, the susceptible cell survives. If not, that susceptible cell dies, which means that the program removes this cell in the simulation’s next steps. If the biological assumption is that persister cells leave the dormant state as soon as their site becomes nontoxic, the computer program follows the approach (b). First, we calculate the difference between the number of susceptible bacteria in the dormant state in the simulations and the one predicted by either the power-law function or the exponential function. If that difference is positive, we randomly chose that number of susceptible cells to leave the dormant state. The program tests if the nearest resistant cell’s distance is less than the total detoxified radius around it. If yes, that cell survives. Otherwise, the program removes this cell from the simulation (the cell dies because the antibiotic is still present). Then we look for all persister cells present in the detoxified area; these cells leave the dormant state, resuming growth. However, if the difference is negative, i.e., if the number of susceptible bacteria in the dormant state in the simulations is lower than the one predicted by the power-law function or the exponential function, we look for all persister cells present in the detoxified area. These cells leave the dormant state, resuming growth.

The population of genetically susceptible cells that resume growth while t <= τ_0_ are, by definition, in the non-persister state, while those susceptible cells that resume growth when t > τ_0_ are persister cells. Each cycle represents a generation time. Here we assume that one generation time is 30 minutes. Time does not flow continuously in the simulations, but rather in intervals of 30 to 30 minutes. Given the division of time into these intervals of 30 minutes, the interval containing τ_0_ has both persister and non-persister bacteria. Using the decay curve of the population of susceptible bacteria, we calculate the percentage of persister and non-persister bacteria in this period. For example, if τ_0_ = 70 mins, the simulation performs 60 mins (two cycles, each representing 30 mins) plus 10 mins decaying as non-persisters, and the remaining 20 mins decaying as persisters. Therefore, 1/3 (=10/30) of the remaining genetically susceptible cells resume growth as non-persisters (i.e., according to the exponential) and 2/3 (=20/30) resume growth as persisters.

At the end of a simulation, we have gathered information on how many persister and non-persister cells generated the population of susceptible cells observed after 24h. Moreover, we can also know how many persister and non-persister cells have survived in each generation.

By performing simulations with several combinations of parameters, we can find those that better explain experimental results.

### Details of the computational model

#### Simulating the medium plate and bacterial cells in the plate

The main procedure was to simulate the experiments performed in Domingues *et al.* (ref. [14]), where susceptible and resistant bacteria were mixed and cultured in agar plates. *Escherichia coli* cells are rod-shaped cells about 2 ፧፧ long and 0.5 ፧፧ diameter, hence occupying a 2-dimensional area of about 1 ፧፧^2^ and a plate has a diameter of 9 cm = 90000 ፧፧. Therefore, we considered that each point in the agar plate, computationally defined by two integers (coordinates *x* and *y*), is the center of a square with an area of 1 ፧፧^2^. The computer program’s first step was to simulate the random distribution of cells in the plate, assigning random coordinates to all cells. Then, we calculated the distances between each susceptible cell and the nearest resistant cell, saving the values in a file.

#### Calculating the number of generations

In each experimental setup, it is possible to calculate how many generations were completed by the resistant population (resistant cells are not affected by the antibiotic). The appropriate mathematical expression is:

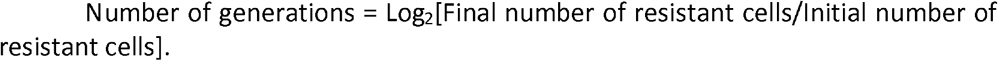

In the simulations, both susceptible and resistant cells belong to the same species and consume the same nutrients. We further assumed that there was no resistance cost, i.e., in the absence of antibiotics, resistant and susceptible cells replicate at the same speed. Therefore, if there were no antibiotics and given that both strains are of the same species, they would complete the same number of generations.

#### Simulating the radial spread of β-lactamase around resistant cells

We simulated the spread of β-lactamase as an expanding circle centered in each resistant bacterium. At any time, these circles represent an antibiotic-free area. According to the Einstein equation for the Brownian motion, the mean displacement of a small particle diffusing in a medium is proportional to the root square of the time elapsed. Therefore, the circle radius grows proportionally to the square root of time, Sqrt(time). Counting the time in bacterial generations, we may express this as R = C.Sqrt(number of generations), where R is the circle’s radius, and C is a constant that depends on the diffusion constant, which may depend on the medium conditions (e.g., the agar concentration). Henceforth, we name this constant C as the “spreading parameter”. Note that in the initial moment (generation 0), the value of R is 0. Therefore, all susceptible bacteria that start dividing at that moment dies.

#### Non-persister versus persister cells and the main parameters

Populations of cells that do not encode for antibiotic resistance die in the presence of bactericidal antibiotics in two phases (Fig 1). In the first phase, between t = 0 and t = τ_0_, the population declines exponentially, i.e., following A_1_·exp(−k_1_·t), where the constants A_1_ and k_1_ are positive. The second phase starts at time t = τ_0_, where the population declines at a slower pace, following a power law or an exponential function, i.e., according to A_2_ .t^፧^, where ፧ is a negative exponent, and A_2_ is a positive constant or according to A_2_·exp(−k_2_·t), where A_2_ and k_2_ are two positive constants and k_2_ < k_1_. All bacteria from this second phase are persister cells. At t = τ_0_, the two mathematical expressions should give the same value, i.e., A_1_·exp(−k_1_·τ_0_) = A_2_. τ_0_^፧^ or A_1_·exp(−k_1_·τ_0_) = A_2_·exp(−k_2_·τ_0_) because the lag time probability distribution is continuous [21]. Moreover, its cumulative probability is equal to 1. Mathematically, this means that the integral of A_1_ ·exp(−k·τ_0_) between t and τ_0_ plus the integral of A_2_. τ_0_^፧^ or A_2_·exp(−k_2_·τ_0_) between τ_0_ and infinity, is equal to 1 [21]. With these two conditions, we can write A_1_ and A_2_ as functions of k_1_, τ_0_ and **β** (assuming that persisters decay according to a power-law) or k_2_ (assuming exponential decay):

For power-law decay: A_1_ = k_1_.exp(k_1_.τ_0_)/R and A_2_ = k_1_/(R.τ_0_^፧^ where R = exp(k_1_.τ_0_) − τ_0_.k_1_/(1+፧)−1
For exponential decay: A_1_ = 1/Q and A_2_ = exp(−(k_1_− k_2_).τ_0_)/Q where Q = (1 − exp(−k_1_.τ_0_))/ k_1_ + exp(−k_1_.τ_0_)/ k_2_

By comparing simulations (this work) with experimental results (obtained in ref. [14]), we can estimate k_1_, τ_0_, and **β** or k_2_.

#### Comparison of results between experiments and simulations

The parameters to adjust were k_1_, τ_0_, ፧ or k_2_, and C. As explained above, the parameters A_1_ and A_2_ depend on k_1_, τ_0_, and ፧ or k_2_. The program ran as many generations as those completed by resistant cells in the experiments performed in Domingues *et al.* [14]. Therefore, the final number of resistant cells should the same both in experiments and simulations.

We ran several simulations by varying the parameters k_1_, τ_0_, ፧ or k_2_, and C, to find the set of parameters that better explain the experimental results found in ref. [14] (Table 1). In these comparisons between experiments and computer simulations, we considered that experiments had an associated experimental error. For instance, agar thickness and other physical conditions of the agar plates that may influence the spreading parameter may constitute a variance source. Furthermore, experiences are also subject to unknown errors. For these reasons, we accept our results to deviate from experimental results. We calculated the lower and upper limits of the intervals according to the following:

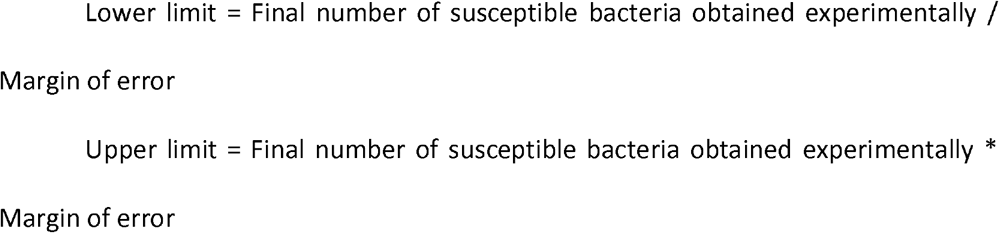

The margins of error tested were 2 and 4.

As explained above, we studied two initial cell densities and three initial frequencies of susceptible to resistant cells. In the simulations, we combined all experimental cases with our parameters. For each combination, we performed three repetitions. In case one repetition result is contained in an interval, we consider that the simulated parameters explain the set experimental results for that margin of error.

## Results

In this work, we took advantage of the experimental results previously obtained in our research group (ref. [14]). The authors spread resistant cells (producers of the detoxifying enzyme β-lactamase) and susceptible cells in a nutrient-rich medium plate with ampicillin (a β-lactam antibiotic), followed by the quantification of susceptible (and resistant) cells after one day. This was done in one of the three frequencies (for a specific initial total density), namely, 99% of susceptible cells and 1% of resistance cells (denominated as 1R:99S), the reverse (99R:1S), and also 50% of each (50R:50S). Resistant cells can produce β-lactamase because they harbor the R1 plasmid encoding the enzyme. This naturally isolated plasmid is transferable by conjugation, so later, we check the impact of conjugation on the survival of susceptible cells.

### The encounter probability of resistant and susceptible cells does not explain the survival of susceptible cells

We started by addressing the hypothesis that surviving susceptible cells are those that were very close to β-lactamase-producing cells, so we analyzed the importance of the encounter probability between resistant and sensitive cells when spread in the agar plate. According to this hypothesis, the probability of encounter between resistant cells and susceptible cells would be the main factor for the survival of susceptible cells. If this was the case, the number of surviving susceptible cells (and their descendants after 24h) should be the same for the 99R:1S and 1R:99S frequencies. The encounter probability of a resistant and a susceptible cell is proportional to 99/100×1/100 (for the case 99R:1S) = 1/100×99/100 (for the case 1R:99S). If the number of surviving susceptible cells was the same, the number of surviving susceptible cells would be similar. However, they differed considerably (Table 1). In the high-density case, the final number of susceptible cells for the frequency 1R:99S was 3.27×10^4^, whereas for 99R:1S was 1.23×10^7^, hence differing by more than three-hundred-fold (Table 1).

The encounter probability for the 50R:50S frequency is proportional to 50/100×50/100. This probability is approximately 25-fold higher than the encounter probability for the 99R:1S and 1R:99S frequencies. Therefore, the above hypothesis predicts that the number of surviving cells in the 50R:50S frequency should be 25 fold higher than in the 99R:1S and 1R:99S frequencies. This prediction is also far from experimental observations (Table 1). For example, for the high-density case, the final number of susceptible cells for the 50R:50S frequency was 1.08×10^6^, which is about 10-fold less, not 25-fold higher than the 1.23×10^7^ cells observed for the 99R:1S frequency (Table 1).

These results suggest that the encounter probability is not an essential factor for the survival of susceptible cells in the indirect resistance phenomenon.

### Persistence is required for susceptible cells survival

After the inoculation of susceptible and resistant cells, the latter replicate for several generations until resources present in the plate are over. The number of generations completed by the resistant cells can be calculated (see the Methods section). Assuming that the resistance cost is negligible and that all susceptible cells start replicating at the same time as resistant cells, we can also estimate how many susceptible cells should have survived when inoculated to explain their final number. Table 3 shows these estimations for the six conditions.

**Table 3.** Estimation of the number of surviving susceptible cells at inoculation time.

| Density | Frequency | Estimated number of surviving susceptible cells at inoculation time |
| --- | --- | --- |
| Low | 1R:99S | $4.85 \times 10^{-5}$ |
| | 50R:50S | $5.33 \times 10^{-6}$ |
| | 99R:1S | $2.27 \times 10^{-4}$ |
| High | 1R:99S | $1.12 \times 10^{-1}$ |
| | 50R:50S | $7.00 \times 10^1$ |
| | 99R:1S | $1.02 \times 10^5$ |

In two cases shown in Table 3 (high density, frequencies 50R:50S and 99R:1S), the estimated number of surviving susceptible cells is higher than one cell, but it was lower than one cell in the other four cases (high density, frequency 1R:99S, and the three frequencies when density was low). These four cases of less than one cell seem unrealistic and need to be understood. A possible explanation is that one or more bacteria have entered the persistence state. In this state, susceptible bacteria can survive in the presence of ampicillin because they are not replicating, and resistant bacteria continue to produce and release β-lactamase into the culturing medium.

The time that each bacterium remains in the persistence state varies from one bacterium to another. When should persistent cells leave the dormant state? We have analyzed four possibilities. The subpopulation of persister cells resumes growth, either according to a power-law or to an exponential-law distribution. For these two cases, dormant cells may or may not resume growth as soon as the medium is nontoxic.

It is impossible to determine the number of persister cells needed to give rise to the final number of susceptible cells observed experimentally. Suppose we observed exactly four surviving susceptible cells at the end of an experiment. We wouldn’t know whether: (i) the four bacteria were in a dormant state all the time; (ii) two cells were in the persistence state most of the time but resumed growth (replicating once) about 30 minutes before the end of the experiment; or (iii) one cell was dormant most of the time but resumed growth about 60 minutes before the end of the experiment. Any of these three scenarios would explain four susceptible cells at the end of the experiment. Increasing the number of final susceptible cells would sharply increase the number of possible scenarios. Therefore, we performed simulations, varying several parameters (more details in the next section), to estimate the number of persister and non-persister cells necessary to explain the experimental number of surviving susceptible cells.

### Simulations to estimate the growth of susceptible cells

We had to consider the spreading of β-lactamase produced by the resistant bacteria and the decline in the susceptible population while exposed to the β-lactam antibiotic. We have seen that this decaying period has two main phases (Fig 1). The non-persistent population decays exponentially until t = τ_0_. At t = τ_0_, only persistent cells survived. They resume growth and die if the antibiotic is still present. In that case, we tested two alternative possibilities for the decay of the persistent population: according to a power-law distribution or according to another exponential distribution.

We used different parameters to describe the population decay: (i) k_1_, the rate constant in the first exponential decay, is the decay rate of the non-persistent population; (ii) τ_0_ is the time from which only persister cells are alive, which is when the probability distribution changes from the exponential decay to the power-law or the second exponential decay; (iii) ***β***, the power-law exponent or k_2_, the rate constant in the second exponential decay. Therefore, in our simulations, we considered these three parameters together (k_1_, τ_0_, and ***β*** or k_1_, τ_0_, and k_2_). We used a fourth parameter (spreading constant) representing the rate increase of the detoxified area – this area increases around each resistant bacterium due to the detoxifying enzyme’s diffusion.

To find the parameters that best fit the experimental results [14], we combined the following parameters: (i) τ_0_ ∈ {20, 30, 50, 60, 70, 80, 90, 100, 110, 120, 130, 150, 200, 250, 300, 350, 400}; (ii) k_1_ ∈ {0.015, 0.020, 0.025, 0.030, 0.040, 0.045, 0.050, 0.055, 0.060, 0.065, 0.070, 0.075, 0.080, 0.090, 0.095, 0.100, 0.200}; (iii) ***β*** ∈ { −1.1, - 1.2, −1.5, −1.7, −1.8, −1.9, −2.0, −2.1, −2.2, −2.3, −2.4, −2.5, −2.7, −2.9, −3.1, −3.3, −3.5} or k_2_ ∈ {0.001, 0.005, 0.010, 0.015, 0.020, 0.025, 0.030, 0.035, 0.040, 0.045, 0.050}, a total of 17^3^ = 4913 combinations (assuming that the persistent population decays according to power-law) or 17*11*11 = 2057 combinations (assuming exponential decay), since k_2_ has to be lower than *k*_l_. For each combination of these parameters, we tested spreading rates C ∈ {0.2, 0.4, 0.6, 0.8, 1, 1.2, 1.4, 1.6, 1.8, 2, 4, 6, 8, 10, 12, 14, 16, 18, 20}. Then, by comparing the computational results with the experimental ones, we obtained a set of parameters that explained the experimental results of the three frequencies (1R:99S, 50R:50S, 99R:1S) in low and high density. We included the possibility of experimental error; that is, we allowed our results to differ from experimental results according to a certain error-margin (see the Methods section).

Assuming that there was no error or that the error-margin is 2, we could not find any combination of parameters (k_1_, τ_0_, ***β***, nor k_1_, τ_0,_ and k_2_) explaining the experimental results. We found six combinations with an error margin of 4 when assuming that persisters decay according to a power-law distribution and two combinations when assuming that persisters decay exponentially (Table 4). The detoxified area’s spreading parameter varied considerably in these combinations, probably due to different experimental conditions (see Discussion).

**Table 4.** The parameters of the simulations that explain the experimental results

| Do persister cells leave<br>the dormant state as<br>soon as the medium<br>becomes detoxified? | Type of<br>decay | $\tau_0$ | $k_1$ | $\beta$ | $k_1$ | Range of spreading<br>parameter |
| --- | --- | --- | --- | --- | --- | --- |
| Yes | Power-law | 20 | 0.07 | -2.3 | NA | 0.4 to 10 |
|  |  | 20 | 0.075 | -2.3 |  | 0.4 to 14 |
|  |  | 20 | 0.08 | -2.2 |  | 0.4 to 10 |
|  |  | 20 | 0.09 | -2.1 |  | 0.4 to 12 |
|  |  | 50 | 0.07 | -2.5 |  | 0.4 to 12 |
|  |  | 60 | 0.07 | -2.2 |  | 0.4 to 14 |
|  | Exponential | 70 | 0.07 | NA | 0.005 | 0.4 to 14 |
|  |  | 80 | 0.065 |  | 0.005 | 0.2 to 10 |
| No | Power-law | 20 | 0.065 | -2.1 | NA | 0.4 to 10 |
|  |  | 20 | 0.07 | -2.1 |  | 0.4 to 10 |
|  |  | 30 | 0.075 | -2.1 |  | 0.4 to 10 |
|  | Exponential | 70 | 0.07 | NA | 0.005 | 0.4 to 14 |
|  |  | 80 | 0.065 |  | 0.005 | 0.2 to 10 |

In the simulations presented until now, persister cells were resuming growth as soon as the medium became detoxified. To understand the impact of this assumption, we repeated the simulations, this time assuming that persister cells stay in the dormant state even when the medium becomes nontoxic. Therefore, in this case, persister cells resume growth stochastically, independently of the antibiotic’s presence in the medium, and, as in the previous case, they do that according to the power-law distribution or the second exponential distribution. We found three combinations with an error margin of 4 if the persistent population decays according to a power-law distribution (Table 4) and one combination if the persistent population decays exponentially (Table 4).

During simulations, we also quantified the number of persistent and non-persistent bacteria throughout generations. Therefore, we can analyze how many susceptible cells in the final population originated from non-persister and how many from persister cells. We have done this analysis for each combination of parameters presented in Table 4. The supporting S1 to S12 Tables show the results of the analyses.

All susceptible bacteria observed at the end of the experiments in the low-density cases descend from persistent bacteria (S1 Table to S12 Table). When the density was high, some non-persistent bacteria also survived in the early generations. As non-persister bacteria duplicate in each generation (contrary to dormant persister cells), they may become strongly represented at the end of the experiment (this is observed for the cases of high density, frequencies 1R:99S and 50R:50S), even if they were a minority of the surviving cells (S1 Table to S12 Table).

When the initial number of β-lactamase-producing cells was too low to detoxify the agar-plate fully, persister cells maintained their state until the end of the experiment (i.e., until 24h later when cells were finally plated in a medium without antibiotic for quantification). Such permanence of bacteria in the persister state occurred when the initial number of β-lactamase-producer cells was low, when the initial total cell density was low (for the three frequencies), or when the initial cell density was high but the initial frequency of β-lactamase-producing cells was low (1R:99S) or intermediate (50R:50S) (S1 Table to S12 Table). In these cases, resistant cells spent all resources before the detoxification of the agar-plate.

### The impact of plasmid transfer in susceptible cells survival is negligible

As explained before, we used experimental data obtained with resistant cells that were encoding the detoxifying enzyme in a transferable (conjugative) plasmid, the R1 plasmid. Therefore, plasmids may move (by replication) into susceptible cells and form transconjugants, here broadly defined as cells that received the R1 plasmid and their descendants. Transconjugants become producers of β-lactamase, hence able to detoxify the environment.

Transconjugants represent a small percentage (between 0% and 1%) of the susceptible cells in the experiments performed with the R1 plasmid by Domingues *et al.* (ref. [14]) (Table 1).

Because we estimated the number of generations completed in the agar-plate, it is possible to assess transconjugants’ impact on indirect resistance. If there are T transconjugants at the end of the experiment, and assuming that the contribution to the detoxification is the highest if all transconjugants formed at the end of the first generation, there would be T/2ng transfer events, where ng is the total number of generations (we are assuming that, at the end of the first generation, resistant cells have already replicated once). In Table 5, we show the number of generations, the final number of transconjugants observed in the agar-plate, the estimated number of plasmid transfers, and the proportion of resistant cells that are transconjugants. In all cases, the proportion of transconjugants among all cells capable of detoxifying the medium was extremely low (0.0011% or less). Therefore, the impact of transconjugants on the detoxification of the medium must have been shallow.

**Table 5.**
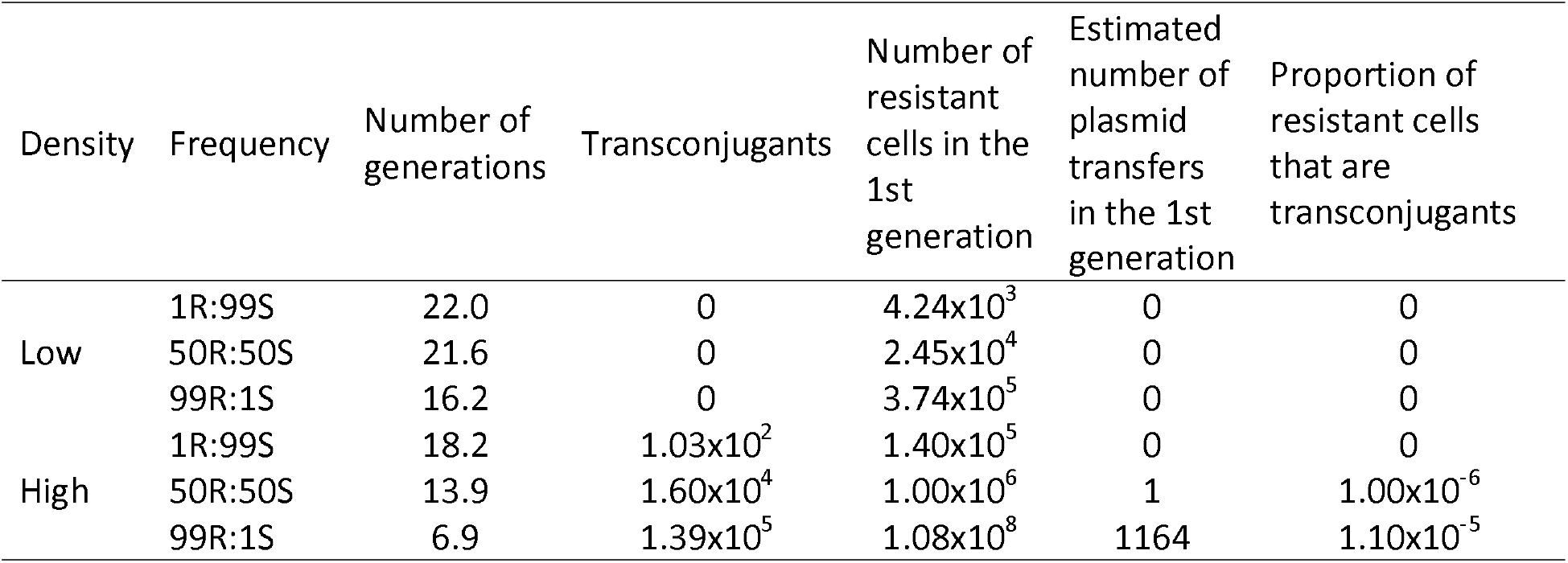
The impact of plasmid transfer to detoxification is low.

| Density | Frequency | Number of generations | Transconjugants | Number of resistant cells in the 1st generation | Estimated number of plasmid transfers in the 1st generation | Proportion of resistant cells that are transconjugants |
| --- | --- | --- | --- | --- | --- | --- |
| Low | 1R:99S | 22.0 | 0 | $4.24 \times 10^3$ | 0 | 0 |
| | 50R:50S | 21.6 | 0 | $2.45 \times 10^4$ | 0 | 0 |
| | 99R:1S | 16.2 | 0 | $3.74 \times 10^5$ | 0 | 0 |
| High | 1R:99S | 18.2 | $1.03 \times 10^2$ | $1.40 \times 10^5$ | 0 | 0 |
| | 50R:50S | 13.9 | $1.60 \times 10^4$ | $1.00 \times 10^6$ | 1 | $1.00 \times 10^{-6}$ |
| | 99R:1S | 6.9 | $1.39 \times 10^5$ | $1.08 \times 10^8$ | 1164 | $1.10 \times 10^{-5}$ |

### Mathematical description of the persistent sub-population and biological implications

Although close to −2, the exponent found by Simsek and Kim (ref. [21]) was - 2.1, as were most exponents found in this study (ranging from −2.5 to −2.1). It seems that the persistent population decays slightly faster than according to 1/t^2^. Therefore, it is relevant to understand how heterogeneous populations should decay. We argue in this section that, if the persistent population is heterogeneous, it should decay according to a distribution close to a power-law but not precisely according to this distribution.

Following the argument by Simsek and Kim (2019), consider a homogeneous population of antibiotic-susceptible cells in the presence of a bactericidal antibiotic. If a non-growing cell rejuvenates (here defined as resuming growth, see below), it dies due to the antibiotic. Therefore, the number of cells still alive at a given time t decreases according to 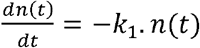, where k_1_ is the rejuvenation probability constant. The solution of this differential equation is *n*(*t*) = *n*(0).*exp*(−*k*_l_. *t*). The rejuvenation probability refers to the number of cells resuming growth in the time interval, which is proportional to the number of cells still alive:

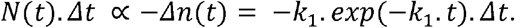

On the other hand, a subpopulation of the cells with various problems in the metabolism, in the cell replication cycle, or even the cell’s response to these problems, stop dividing for some time [20,21]. Each bacterium may have a different issue from a big group of possible problems. Therefore, these bacteria should present a wide range of rejuvenation constants [21]. This bacterial population is heterogeneous, with many different k constants. The number of cells resuming growth at a particular time t, *N*(*t*), is proportional to:

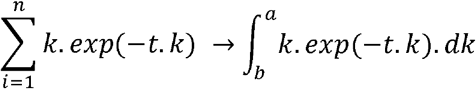

If the population decays between time *τ_0_* and *t_Max_*, then the integral’s limits are *a > 1/τ_0_* and *b < 1/t_Max_*. This lower limit b is close to zero because *t_Max_* is high - the persistent population endures a long time [21].

Note that, until now, we only know that the upper limit of the integral, *a*, has to be higher than *1*/τ_0_. We now argue that this upper limit has to be lower than k_1_. This limit arises from the fact that cells in this heterogeneous sub-population rejuvenate later than the non-persister cells - their rejuvenation constant should be lower than that of the non-persister cells. Therefore, the integral becomes:

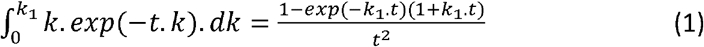

where k_1_ is the rejuvenation constant of the non-persister population.

In general, 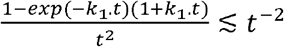.

Experimental results from ref. [21] have shown that the persistent population starts decaying after about 93 min and k_1_ is about 0.063 min^−1^. Therefore, k_1_.t ⋍ 5.859 or higher and increases in time, so the numerator in Equation 1 is 0.98 (that is 1-*exp*(−*k*_1_.*t*)(1+*k*_1_.*t*) = 0.98). Therefore, in general 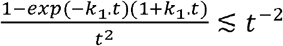. This result may explain why Simsek and Kim (ref. [21]) derived an exponent from their experiments of −2.1, which is slightly lower than their theoretical prediction of −2. However, when t increases, the numerator of Equation 1 converges to 1, which means that the power-law t^β^ should converge to t^−2^ when *t* increases.

Likewise, our results for the power-law decay (Table 4) suggest that the persistent population starts decaying after about 20 to 60 min, and k_1_ is between 0.065 and 0.09 min^−1^. Therefore, k_1_.t ⋍ between 1.3 and 4.2 and increases in time (because t increases), so the numerator in Equation 1 is between 0.37 and 0.92. Again, our results suggest that 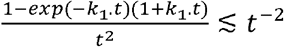 and also explains why we obtained exponents slightly lower than −2.

In this work, we show the involvement of persistent cells during the process of indirect resistance, even in short-time experiments of 24h (like the ones performed in ref. [14]), and that, most likely, persister cells decay according to t^β^ where β is slightly lower than −2.

## Discussion

To understand persisters’ behavior, we started by asking whether they were responsible for the survival of susceptible cells in the context of indirect resistance. For that, we carried out simulations to mimic the experiments that we have performed in a previous work where we spread a mixture of susceptible and β-lactamase-producing cells in agar-plates supplemented with a β-lactam antibiotic [14]. We simulated the behavior of persister cells in four different ways: (i) in the presence of a bactericidal antibiotic, the persistent population decays according to an exponential-law versus according to a power-law; (ii) persister cells leave the dormant state as soon as the medium becomes detoxified *versus* independently of the medium detoxification, hence merely according to the probability mentioned above. Our simulations suggest that persister cells and their descendants were a part, or even all, of the surviving susceptible population, irrespectively of the four alternative behavior models of the persister cells implemented in the simulations. Persisters were the only survivors in the indirect resistance phenomenon when the initial cell density was low.

Given persistent cells’ involvement, we used the results to go more in-depth and understand their nature. Arguably, the prevalent view is that persistence is an evolved characteristic. If genetically encoded, the expectation would be that the persistent population is homogeneous and decays exponentially [25]. Instead, a few recent works have proposed that persistence is an accidental consequence of inadvertent cell problems and errors [19,20]. In this case, the persistent populations should be heterogeneous because cells would have different reasons for showing low metabolism, and the consequent theoretical prediction is that the persistent population should decay, not exponentially, but according to a power-law with the exponent of −2 [21] or slightly lower than −2 (this paper).

The exponential decays are direct consequences of first-order kinetics. The exponential declines occur in various situations, from radioactive decay to the drop of atmospheric pressure with increasing height above sea level. And, of course, the non-persister bacterial population also decays exponentially in time because the bacterial population is large, homogenous, and the law of large numbers holds. There is no theoretical prediction for the decay rate (so far) if the persistent population declines exponentially. Our simulations show that an exponential decline of persisters is possible only for shallow values of the decay constant - this allows the survival of persister cells for several hours in the experiments. However, Simsek and Kim (ref. [21]) were able to mathematically predict the exponent in the power-law case, namely that it should be −2. Likewise, we found exponents close to −2 in the simulations where we assume that persisters’ decay follows a power-law (Table 4).

It is relevant to emphasize that, despite the similarity of the exponent values found here (based on the experiments from ref. [14]) and in the Simsek and Kim’ study [21], the experimental methods of these two studies were significantly different. While Simsek and Kim [21] studied the decay of the susceptible population in a liquid and well-mixed medium without resistant cells, the experiments simulated here (based in ref. [14]) were performed in agar-plates where some susceptible cells die due to the antibiotic and others survive thanks to persistence or the effect of indirect resistance. The similarity of the exponents found with two different experimental methods, with the one predicted theoretically [21], suggests that the persistent population indeed decays according to power-law.

We evaluated the impact of persister cells resuming growth as soon as the medium is nontoxic versus resuming growth stochastically, independently of the antibiotic’s presence in the medium. We found reasonable sets of parameters using both behavioral models. Therefore, strictly speaking, we could not conclude whether persister cells leave the dormant state and resume growth when the medium is nontoxic. However, returning to growth immediately after detoxification implies a sensing mechanism, suggesting that persistence is an evolved mechanism, not the result of inadvertent metabolic and cell replication problems. In that way, it is contradictory to assume simultaneously that persistent cells leave the dormant state as soon as the medium is nontoxic and that the persistent population decays according to power-law. These were the assumptions leading to the first six lines of Table 4. Consequently, we should discard the exponents shown in this part of Table 4. We conclude that the exponent value of the power-law t^β^ is β = −2.1 (lines 9 to 11 of Table 4). This value is the same exponent experimentally measured by Simsek and Kim (ref. [21]).

The coincidence of the exponent values in this work and Simsek and Kim (ref. [21]) work is impressive, but there should be an explanation for the discrepancy from the theoretical prediction of −2. We have shown that heterogeneous populations should decay according to 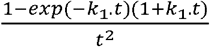, which is close to but slightly lower than 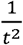. Such discrepancy may explain why our simulations and Simsek and Kim’s experiments point to exponents slightly lower than −2. Both theoretical predictions assumed that several sub-populations of cells constitute the persistent population. The difference between the two theoretical predictions is that our derivation assumes that no hypothetic subpopulations are decaying faster than non-persister cells. This assumption implies fewer cells alive in the persistent state than predicted before [21]. As time passes, the two mathematical predictions converge because even if we were including the subpopulations decaying faster than the non-persistent population, those cells would already be dead.

Given that we simulated bacteria in the agar-plate, we had to consider the radial spreading of β-lactamase around their producers (resistant cells) and the subsequent decrease in antibiotic concentration. The system has some complexity because, in some simulations, the initial number of resistant cells can be high and because there is undoubtedly diffusion of β-lactamase from each resistant bacterium and of the antibiotic towards each resistant bacterium. It is even possible that the detoxifying area increases as a diffusion wave. Moreover, resistant cells duplicate every half an hour, probably increasing the β-lactamase enzyme production outwards the resistant colony. Therefore, we assumed that the detoxified area increases monotonically in time. Future studies should scrutinize the relevance of this assumption. We had to consider a wide range of values for the detoxified area’s speed of increase to fit the experimental results. This range may have several causes. For example, although the agar concentration was the same in all experiments, some plates could be more dried than others, eventually facilitating or hampering the detoxifying enzyme molecules and antibiotic molecules’ movement.

Table 5 shows that transconjugants’ participation in the detoxification of the agar-plate must have been low. This result agrees with previous works showing that the transfer rate of the R1 plasmid is low [14,22,24,26,27].

Several resistance determinants, including genes and chromosomal mutations, are responsible for the burden of antibiotic resistance. This burden is tremendous. Just in the European Economic Area, antibiotic resistance is responsible for 33000 deaths/year and 874000 disability-adjusted life-years [28]. Unfortunately, to survive bactericidal antibiotics, bacteria do not even need to harbor resistance determinants. Susceptible bacteria may rely on indirect resistance and bacterial persistence, as we have seen. Therefore, this work’s conclusions that persistence is often involved in indirect resistance and that persister cells seem to decay according to a power-law are worrying.

The power-law distribution has a long tail, which means that, at least theoretically, some susceptible bacteria may survive for several weeks, eventually after the end of antibiotic uptake by the patient. Long-lived persisters may dictate treatments’ failure because some of these cells may leave the dormant state and reinitiate their pathogenic effects. This risk goes in line with the reports on persistence being a significant cause for recurrent and chronic infections, dictating the patients’ disease progression and outcome [29,30]

In conclusion, this work supports the hypothesis that the persistent population decays according to a power-law with an exponent close to −2. As Simsek and Kim (ref. [21]) argued, such power-law decay means that persistence is the consequence of accidental problems involving replication and metabolism, instead of being an evolved character (see also [19,20]). If confirmed, the implication is that persistence is maladaptive, despite its frequent dramatic medical consequences. A strategy to find anti-persistent drugs should perhaps be different if persisters are moribund cells versus the result of an evolved genetic program.

## Acknowledgments

We thank Octávio Paulo and Francisco Pina-Martins for access to the server.

## Supporting information captions

S1 Table. **Persister and non-persister cells that originated the final susceptible population considering τ_0_ = 20, *k*_1_ = 0.07, *β* = −2.3.**

Results of simulations when we assumed that the persister population decays according to a power-law and that persister cells leave the dormant state as soon as the medium becomes detoxified.

S2 Table. **Persister and non-persister cells that originated the final susceptible population considering τ_0_ = 20, *k*_1_ = 0.075, *β* = −2.3.**

Results of simulations when we assumed that the persister population decays according to a power-law and that persister cells leave the dormant state as soon as the medium becomes detoxified.

S3 Table. **Persister and non-persister cells that originated the final susceptible population considering τ_0_ = 20, *k*_1_ = 0.08, *β* = −2.2.**

Results of simulations when we assumed that the persister population decays according to a power-law and that persister cells leave the dormant state as soon as the medium becomes detoxified.

S4 Table. **Persister and non-persister cells that originated the final susceptible population considering τ_0_ = 20, *k*_1_ = 0.09, *β* = −2.1.**

Results of simulations when we assumed that the persister population decays according to a power-law and that persister cells leave the dormant state as soon as the medium becomes detoxified.

S5 Table. **Persister and non-persister cells that originated the final susceptible population considering τ_0_ = 50, *k*_1_ = 0.07, *β* = −2.5.**

Results of simulations when we assumed that the persister population decays according to a power-law and that persister cells leave the dormant state as soon as the medium becomes detoxified.

S6 Table. **Persister and non-persister cells that originated the final susceptible population considering τ_0_ = 60, *k*_1_ = 0.07, *β* = −2.2.**

Results of simulations when we assumed that the persister population decays according to a power-law and that persister cells leave the dormant state as soon as the medium becomes detoxified.

S7 Table. **Persister and non-persister cells that originated the final susceptible population considering τ_0_ = 70, *k*_1_ = 0.07, *k*_2_ = 0.005.**

Results of simulations when we assumed that the persister population decays according to an exponential-law and that persister cells leave the dormant state as soon as the medium becomes detoxified.

S8 Table. **Persister and non-persister cells that originated the final susceptible population considering τ_0_ = 80, *k*_1_ = 0.065, *k*_2_ = 0.005.**

Results of simulations when we assumed that the persister population decays according to an exponential-law and that persister cells leave the dormant state as soon as the medium becomes detoxified.

S9 Table. **Persister and non-persister cells that originated the final susceptible population considering τ_0_ = 20, *k*_1_ = 0.065, *β* = −2.1.**

Results of simulations when we assumed that the persister population decays according to a power-law and that persister cells do not leave the dormant state as soon as the medium becomes detoxified.

S10 Table. **Persister and non-persister cells that originated the final susceptible population considering τ_0_ = 20, *k*_1_ = 0.07, *β* = −2.1.**

Results of simulations when we assumed that the persister population decays according to a power-law and that persister cells do not leave the dormant state as soon as the medium becomes detoxified.

S11 Table. **Persister and non-persister cells that originated the final susceptible population considering τ_0_ = 30, *k*_1_ = 0.075, *β* = −2.1.**

Results of simulations when we assumed that the persister population decays according to a power-law and that persister cells do not leave the dormant state as soon as the medium becomes detoxified.

S12 Table. **Persister and non-persister cells that originated the final susceptible population considering τ_0_ = 80, *k*_1_ = 0.065, *k*_2_ = 0.005.**

Results of simulations when we assumed that the persister population decays according to an exponential-law and that persister cells do not leave the dormant state as soon as the medium becomes detoxified.

