## Supplementary material for "The power of dying slowly - persistence as unintentional dormancy": S1 Table

| S1 Table - Persister and non-persister cells that originated the final susceptible population considering τ_0_ = 20, $\boldsymbol{k}_{\boldsymbol{1}}$ = 0.07, $\boldsymbol{\beta}$ = -2.3* | | | | | |
| --- | --- | --- | --- | --- | --- |
| **Density** | **Frequency** | **Persister bacteria (%)** | **Total non-persister survivors (without considering duplications)** | **Total persister survivors (without considering duplications)** | **Total dormant cells at the end of the simulations** |
| Low | 1R:99S | 100 | 0 | 565 | 565 |
|  | 50R:50S | 100 | 0 | 27 | 27 |
|  | 99R:1S | 100 | 0 | 8 | 8 |
| High | 1R:99S | 38 | 1 | 9823 | 9819 |
|  | 50R:50S | 48 | 316 | 3190 | 2240 |
|  | 99R:1S | 36 | 66156 | 64041 | 0 |

* We assumed that the persister population decays according to a power-law and that persister cells leave the dormant state as soon as the medium becomes detoxified
