## Supplementary material for "The power of dying slowly - persistence as unintentional dormancy": S3 Table

| S3 Table - Persister and non-persister cells that originated the final susceptible population considering τ_0_ = 20, $\boldsymbol{k}_{\boldsymbol{1}}$ = 0.08, $\boldsymbol{\beta}$ = -2.2* | | | | | |
| --- | --- | --- | --- | --- | --- |
| **Density** | **Frequency** | **Persister bacteria (%)** | **Total non-persister survivors (without considering duplications)** | **Total persister survivors (without considering duplications)** | **Total dormant cells at the end of the simulations** |
| Low | 1R:99S | 100 | 0 | 926 | 926 |
|  | 50R:50S | 100 | 0 | 44 | 44 |
|  | 99R:1S | 100 | 0 | 21 | 21 |
| High | 1R:99S | 40 | 1 | 22905 | 22887 |
|  | 50R:50S | 31 | 455 | 9039 | 6998 |
|  | 99R:1S | 8 | 109208 | 64753 | 0 |

* We assumed that the persister population decays according to a power-law and that persister cells leave the dormant state as soon as the medium becomes detoxified
