## Supplementary material for "The power of dying slowly - persistence as unintentional dormancy": S4 Table

| S4 Table - Persister and non-persister cells that originated the final susceptible population considering τ_0_ = 20, $\boldsymbol{k}_{\boldsymbol{1}}$= 0.09, $\boldsymbol{\beta}$ = -2.1* | | | | | |
| --- | --- | --- | --- | --- | --- |
| **Density** | **Frequency** | **Persister bacteria (%)** | **Total non-persister survivors (without considering duplications)** | **Total persister survivors (without considering duplications)** | **Total dormant cells at the end of the simulations** |
| Low | 1R:99S | 100 | 0 | 689 | 689 |
|  | 50R:50S | 100 | 0 | 33 | 33 |
|  | 99R:1S | 100 | 0 | 16 | 16 |
| High | 1R:99S | 15 | 2 | 17027 | 17026 |
|  | 50R:50S | 22 | 489 | 6717 | 5206 |
|  | 99R:1S | 6 | 114308 | 48121 | 0 |

* We assumed that the persister population decays according to a power-law and that persister cells leave the dormant state as soon as the medium becomes detoxified
