## Supplementary material for "The power of dying slowly - persistence as unintentional dormancy": S5 Table

| S5 Table - Persister and non-persister cells that originated the final susceptible population considering τ_0_ = 50, $\boldsymbol{k}_{\boldsymbol{1}}$ = 0.07, $\boldsymbol{\beta}$ = -2.5* | | | | | |
| --- | --- | --- | --- | --- | --- |
| **Density** | **Frequency** | **Persister bacteria (%)** | **Total non-persister survivors (without considering duplications)** | **Total persister survivors (without considering duplications)** | **Total dormant cells at the end of the simulations** |
| Low | 1R:99S | 100 | 0 | 286 | 286 |
|  | 50R:50S | 100 | 0 | 14 | 14 |
|  | 99R:1S | 100 | 0 | 5 | 4 |
| High | 1R:99S | 4 | 2 | 5163 | 5162 |
|  | 50R:50S | 18 | 640 | 1997 | 1230 |
|  | 99R:1S | 6 | 109169 | 24845 | 0 |

* We assumed that the persister population decays according to a power-law and that persister cells leave the dormant state as soon as the medium becomes detoxified
