## Supplementary material for "The power of dying slowly - persistence as unintentional dormancy": S6 Table

| S6 Table - Persister and non-persister cells that originated the final susceptible population considering τ_0_ = 60, $\boldsymbol{k}_{\boldsymbol{1}}$ = 0.07, $\boldsymbol{\beta}$ = -2.2* | | | | | |
| --- | --- | --- | --- | --- | --- |
| **Density** | **Frequency** | **Persister bacteria (%)** | **Total non-persister survivors (without considering duplications)** | **Total persister survivors (without considering duplications)** | **Total dormant cells at the end of the simulations** |
| Low | 1R:99S | 100 | 0 | 584 | 584 |
|  | 50R:50S | 100 | 0 | 28 | 28 |
|  | 99R:1S | 100 | 0 | 8 | 8 |
| High | 1R:99S | 7 | 2 | 9959 | 9957 |
|  | 50R:50S | 11 | 716 | 3044 | 2212 |
|  | 99R:1S | 2 | 114588 | 19390 | 0 |

* We assumed that the persister population decays according to a power-law and that persister cells leave the dormant state as soon as the medium becomes detoxified
