## Supplementary material for "The power of dying slowly - persistence as unintentional dormancy": S7 Table

| S7 Table - Persister and non-persister cells that originated the final susceptible population considering τ_0_ = 70, $\boldsymbol{k}_{\boldsymbol{1}}$ = 0.07, $\boldsymbol{k}_{\boldsymbol{2}}$ = 0.005* | | | | | |
| --- | --- | --- | --- | --- | --- |
| **Density** | **Frequency** | **Persister bacteria (%)** | **Total non-persister survivors (without considering duplications)** | **Total persister survivors (without considering duplications)** | **Total dormant cells at the end of the simulations** |
| Low | 1R:99S | 100 | 0 | 779 | 779 |
|  | 50R:50S | 100 | 0 | 37 | 37 |
|  | 99R:1S | 100 | 0 | 11 | 11 |
| High | 1R:99S | 40 | 1 | 13290 | 13284 |
|  | 50R:50S | 50 | 308 | 4031 | 2961 |
|  | 99R:1S | 37 | 64250 | 67201 | 0 |

***** We assumed that the persister population decays according to an exponential-law and that persister cells leave the dormant state as soon as the medium becomes detoxified
