## Supplementary material for "The power of dying slowly - persistence as unintentional dormancy": S8Table

| S8 Table - Persister and non-persister cells that originated the final susceptible population considering τ_0_ = 80, $\boldsymbol{k}_{\boldsymbol{1}}$ = 0.065, $\boldsymbol{k}_{\boldsymbol{2}}$ = 0.005* | | | | | |
| --- | --- | --- | --- | --- | --- |
| **Density** | **Frequency** | **Persister bacteria (%)** | **Total non-persister survivors (without considering duplications)** | **Total persister survivors (without considering duplications)** | **Total dormant cells at the end of the simulations** |
| Low | 1R:99S | 100 | 0 | 1077 | 1077 |
|  | 50R:50S | 100 | 0 | 52 | 52 |
|  | 99R:1S | 100 | 0 | 14 | 14 |
| High | 1R:99S | 42 | 1 | 18027 | 18020 |
|  | 50R:50S | 53 | 228 | 4888 | 3929 |
|  | 99R:1S | 38 | 64316 | 71322 | 0 |

***** We assumed that the persister population decays according to an exponential-law and that persister cells leave the dormant state as soon as the medium becomes detoxified
