## Supplementary material for "The power of dying slowly - persistence as unintentional dormancy": S9 Table

| S9 Table - Persister and non-persister cells that originated the final susceptible population considering τ_0_ = 20, $\boldsymbol{k}_{\boldsymbol{1}}$ = 0.065, $\boldsymbol{\beta}$ = -2.1* | | | | | |
| --- | --- | --- | --- | --- | --- |
| **Density** | **Frequency** | **Persister bacteria (%)** | **Total non-persister survivors (without considering duplications)** | **Total persister survivors (without considering duplications)** | **Total dormant cells at the end of the simulations** |
| Low | 1R:99S | 100 | 0 | 1350 | 1350 |
|  | 50R:50S | 100 | 0 | 65 | 65 |
|  | 99R:1S | 100 | 0 | 18 | 18 |
| High | 1R:99S | 42 | 1 | 22598 | 22594 |
|  | 50R:50S | 38 | 362 | 6030 | 4945 |
|  | 99R:1S | 30 | 74904 | 91866 | 17420 |

*We assumed that the persister population decays according to a power-law and that persister cells do not leave the dormant state as soon as the medium becomes detoxified
