## Supplementary material for "The power of dying slowly - persistence as unintentional dormancy": S11 Table

| S11 Table - Persister and non-persister cells that originated the final susceptible population considering τ_0_ = 30, $\boldsymbol{k}_{\boldsymbol{1}}$ = 0.075, $\boldsymbol{\beta}$ = -2.1 | | | | | |
| --- | --- | --- | --- | --- | --- |
| **Density** | **Frequency** | **Persister bacteria (%)** | **Total non-persister survivors (without considering duplications)** | **Total persister survivors (without considering duplications)** | **Total dormant cells at the end of the simulations** |
| Low | 1R:99S | 100 | 0 | 1335 | 1335 |
|  | 50R:50S | 100 | 0 | 64 | 64 |
|  | 99R:1S | 100 | 0 | 17 | 17 |
| High | 1R:99S | 14 | 2 | 22337 | 22334 |
|  | 50R:50S | 17 | 556 | 5884 | 4888 |
|  | 99R:1S | 7 | 108127 | 53878 | 17219 |

* We assumed that the persister population decays according to a power-law and that persister cells do not leave the dormant state as soon as the medium becomes detoxified
